# Mpox virus (MPXV) vertical transmission and fetal demise in a pregnant rhesus macaque model

**DOI:** 10.1101/2024.05.29.596240

**Authors:** Nicholas P. Krabbe, Ann M. Mitzey, Saswati Bhattacharya, Elaina R. Razo, Xiankun Zeng, Nell Bekiares, Amy Moy, Amy Kamholz, Julie A. Karl, Gregory Daggett, Grace VanSleet, Terry Morgan, Saverio V. Capuano, Heather A. Simmons, Puja Basu, Andrea M. Weiler, David H. O’Connor, Thomas C. Friedrich, Thaddeus G. Golos, Emma L. Mohr

## Abstract

Infection with clade I Mpox virus (MPXV) results in adverse pregnancy outcomes, yet the potential for vertical transmission resulting in fetal harm with clade IIb MPXV, the clade that is currently circulating in the Western Hemisphere, remains unknown. We established a rhesus macaque model of vertical MPXV transmission with early gestation inoculation. Three pregnant rhesus macaques were inoculated intradermally with 1.5 × 10^5 plaque forming units (PFU) of clade IIb MPXV near gestational day (GD) 30 and animals were monitored for viremia and maternal and fetal well-being. Animals were euthanized to collect tissues at 5, 14, or 25 days post-inoculation (dpi). Tissues were evaluated for viral DNA (vDNA) loads, infectious virus titers, histopathology, MPXV mRNA and protein localization, as well as MPXV protein co-localization with placental cells including, Hofbauer cells, mesenchymal stromal cells, endothelial cells, and trophoblasts. vDNA was detected in maternal blood and skin lesions by 5 dpi. Lack of fetal heartbeat was observed at 14 or 25 dpi for two dams indicating fetal demise; the third dam developed significant vaginal bleeding at 5 dpi and was deemed an impending miscarriage. vDNA was detected in placental and fetal tissue in both fetal demise cases. MPXV localized to placental villi by ISH and IHC. Clade IIb MPXV infection in pregnant rhesus macaques results in vertical transmission to the fetus and adverse pregnancy outcomes, like clade I MPXV. Further studies are needed to determine whether antiviral therapy with tecovirimat will prevent vertical transmission and improve pregnancy outcomes.

**One Sentence Summary:** Clade IIb Mpox virus infection of pregnant rhesus macaques results in vertical transmission from mother to fetus and adverse pregnancy outcomes.

## INTRODUCTION

Mpox (formerly monkeypox) virus (MPXV) is a serious health concern that recently spread around the world, resulting in over 94,000 infections globally (*1*). Some MPXV infections cause fetal demise, particularly with clade I MPXV, which is endemic in Central Africa (reviewed in (*2*)). It is unclear whether clade IIb MPXV, which is circulating worldwide including the United States, is also vertically transmitted and causes adverse pregnancy outcomes.

In 2022, an outbreak of mpox, resulting from infection with clade IIb MPXV, spread rapidly across Europe, the Americas, and a total of 110 countries worldwide. MPXV infections have historically been reported in West as well as East and Central Africa (*3*), with clade I MPXV reported in Central and East Africa and clade II in West Africa (*4*) although there is concurrent circulation of both clades in some countries (*5*). The United States has reported over 32,000 cases of MPXV infection and 58 deaths as of March 2024 (*1*). Although the number of cases peaked in 2022 and numbers have decreased since then, new cases are reported every week within the United States (*1*). This highlights the unpredictability of MPXV infection, and the need to better define disease outcomes in vulnerable populations.

Clade I MPXV infection during pregnancy is associated with high (up to 75%) rates of fetal demise (*6*). Fetal demise may be due to vertical transmission because fetuses developed diffuse cutaneous maculopapular lesions, hepatomegaly, and hydrops fetalis. Additionally, MPXV DNA was identified in the placenta and viral proteins were identified in the chorionic villi and fetal skin by immunohistochemistry (*6, 7*). The number of women impacted by clade I MPXV infection during pregnancy is unknown, but the risk to pregnant persons is serious and ongoing because there have been over 40 outbreaks in the Central African Republic alone since 2018 (*8*) and more than 19,000 cases in the Democratic Republic of Congo since 2023 (*9*). The outcomes of clade IIb MPXV infection during pregnancy are not well described, with few publications characterizing the outcomes of pregnant persons in the United States (*10, 11*). Oakley et al. described a cohort of pregnancies where MPXV infection occurred at some point during pregnancy (*10*). Of the 3 pregnancies with reported outcomes, 2 had full-term deliveries without complications and one had a spontaneous abortion at 11 weeks gestation (*10*). A recent case report described MPXV virus (presumed to be clade IIb based on location in the United States) infection at 31 weeks gestation that resulted in a full-term delivery with no apparent infant infection (*11*). This case was treated with the antiviral medication tecovirimat and while no side effects were reported, it is unclear whether any adverse events from tecovirimat were evaluated (*11*). Determining whether clade IIb MPXV infection during pregnancy increases the risk of fetal demise and vertical transmission is critical to guide risk assessments and treatment recommendations during pregnancy (*2, 12–14*).

We developed a rhesus macaque model of prenatal MPXV infection to determine whether clade IIb MPXV can be vertically transmitted and determine whether fetal demise occurs. We modeled human infection as much as possible, using an intradermal inoculation route, a low passage clade IIb MPXV isolate, and a dose similar to what may be transmitted by skin-to-skin contact in this translational model. We found that clade IIb MPXV infection in the first trimester led to maternal viremia, the development of maternal skin lesions, vertical transmission as evidenced by the presence of MPXV in maternal-fetal interface tissues, amniotic fluid, and fetal tissues, and fetal demise in 2 of the 3 cases.

## RESULTS

### Intradermal inoculation results in systemic MPXV infection

We inoculated three pregnant Indian-origin rhesus macaques (Macaca mulatta) intradermally around gestational day (GD) 30 with clade IIb MPXV derived from a human isolate. To ensure that the challenge stock accurately reflected the virus obtained from BEI Resources, the stock was sequenced using long-read Oxford Nanopore sequencing for accurate reconstruction of repetitive sequences in the MPXV genome. After read mapping to the expected MPXV reference sequence (GenBank accession: ON563414.3), a mean of 281x coverage (163x standard deviation) was obtained, with no regions of sequence coverage <20x. When considering variants detected at ≥25% frequency, there were only three single-nucleotide polymorphisms (SNPs) relative to the reference sequence. Three insertions of 128bp, 128bp, and 342bp were found along with several small deletions. Taken together, the sequence of the challenge stock retained high fidelity to the parental virus.

All dams developed MPXV viremia by 2-5 days post inoculation (Fig. 1A) and skin lesions by 4-7 days post-inoculation (Fig. 1B). Skin lesion swabs had high viral loads, with one animal demonstrating increasing skin lesion viral loads following inoculation (ID 101), and the other with consistently high skin lesion viral loads (ID 102) (Fig. 1C). Skin lesions first appeared as erythematous macules which progressed to papules, pustules, and ulcerations with multiple skin lesion stages present simultaneously in each dam in the two fetal demise cases (Fig. S1). The dam with an impending miscarriage had multiple intraepidermal pustules near the injection sites but did not have skin lesions sampled prior to euthanasia, so no viral loads were available.

**Fig 1.**
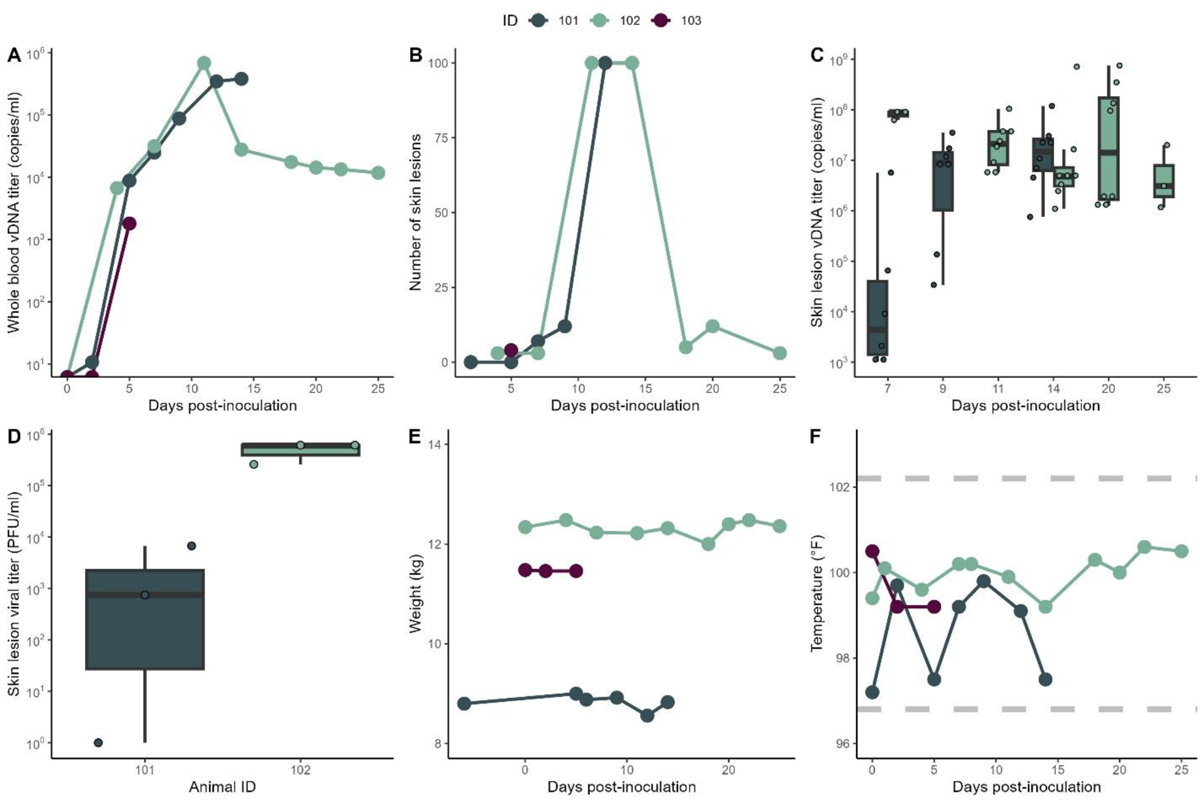
Maternal systemic MPXV infection. (**A**) Whole blood MPXV DNA viral loads. (**B**) Skin lesion counts. (**C**) MPXV DNA viral loads in eluates from swabbed skin lesions. (**D**) Infectious virus titers in eluates from swabbed skin lesions measured by plaque assay at 7 days post-inoculation. (**E**) Maternal weight. (**F**) Maternal temperature, with the normal temperature range shown between the dashed lines.

Eluates of skin lesion swabs taken at 7 days post-inoculation had replication-competent virus present (except for one lesion from ID 101), with an average titer of 2.5×10^5^ plaque forming units (PFU)/mL between both dams (Fig. 1D). This average titer was similar to our intradermal inoculation dose of 1.5 x10^5^ PFU. Skin lesions consisted of intra-epidermal pustules, ballooning degeneration of keratinocytes, and neutrophilic dermatitis and vasculitis with multifocal ulceration as well as viral DNA and proteins identified by ISH and IHC (Fig. S2). None of the dams developed weight loss (Fig. 1E), although it should be noted that all animals received supplementation with oral gavage feeds because of decreased appetite noticed by veterinary services. No fever was detected from inoculation at any time (Fig. 1F).

### MPXV is vertically transmitted

Fetal demise was identified in two pregnancies and an impending miscarriage was identified in a third pregnancy. One fetal demise was identified at 14 days post-inoculation (ID 101), another demise at 25 days post-inoculation (ID 102), and an impending miscarriage at 5 days post-inoculation (ID 103). Each demise was identified by the lack of fetal heartbeat present on ultrasound. The impending miscarriage was characterized by significant vaginal bleeding which worsened with abdominal palpation and was verified by the visualization of a large amount of intrauterine fluid between the internal cervical os and amniotic sac on ultrasound. Because active intrauterine hemorrhage, abruption, and imminent fetal loss were suspected, euthanasia was recommended by a veterinarian. All dams were euthanized the day of fetal demise or impending fetal demise with subsequent necropsy examination.

There were high MPXV DNA viral loads in multiple fetal samples in both fetal demise cases, including the brain, eye, heart, kidney, liver, lung, skin, umbilical cord, and cerebrospinal fluid (Fig. 2). In contrast, there was no MPXV DNA isolated in the fetal tissues from the impending miscarriage case, at 5 days post-inoculation. The two fetal demise cases had higher viral DNA loads in fetal cerebrospinal fluid than in the maternal-fetal interface (MFI) fluids (amniotic fluid, chorionic fluid), or maternal fluids (cerebrospinal fluid, urine, vaginal secretions) (Fig. 2).

**Fig 2.**
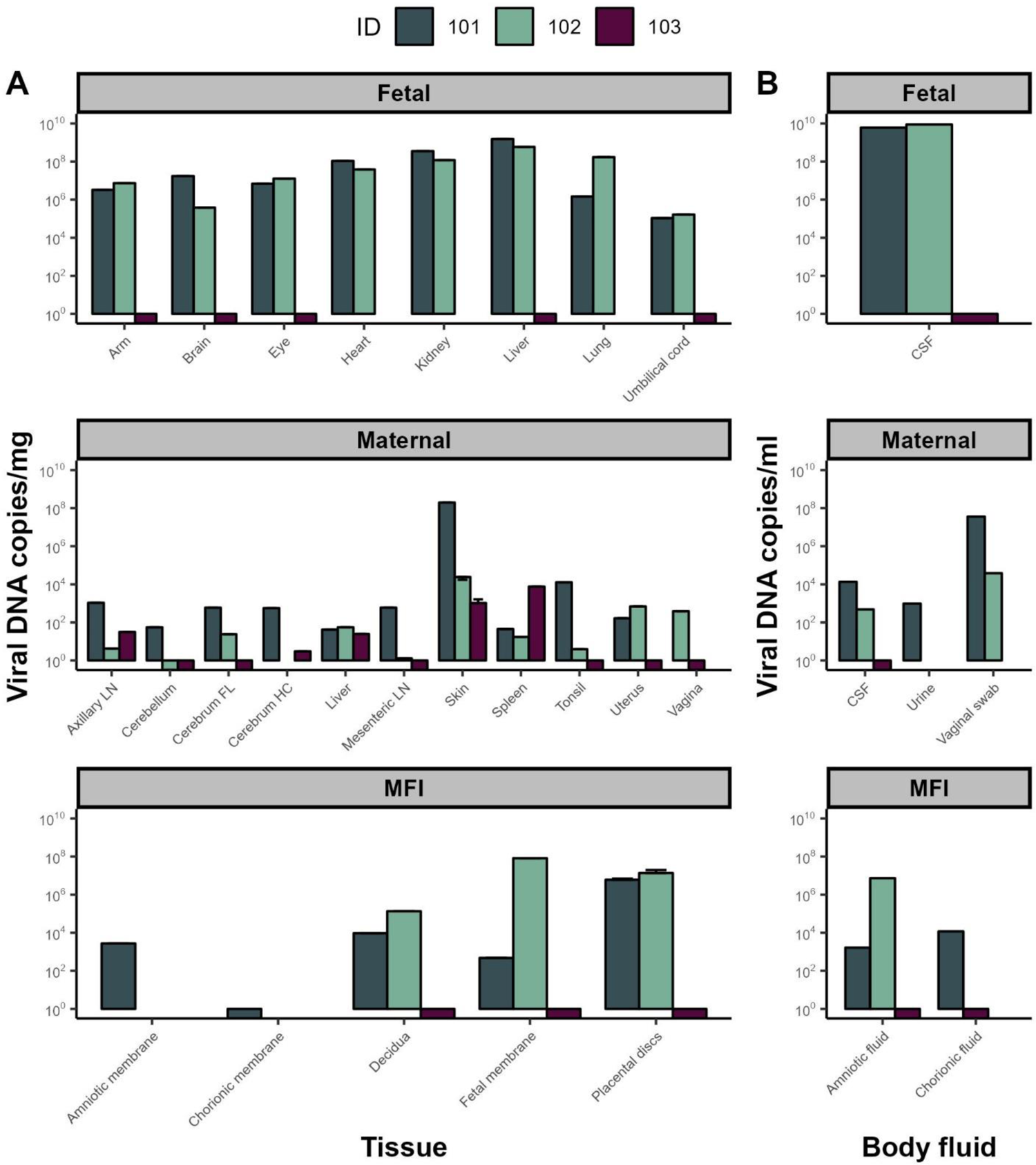
MPXV tissue and body fluid viral loads. MPXV viral loads were measured at the time of necropsy in fetal, maternal, and maternal-fetal interface (MFI) tissues (**A**) and body fluids (**B**). Cerebrospinal fluid (CSF), lymph node (LN), frontal lobe (FL), hippocampus (HC). The average viral load with standard deviation is shown for placental discs and skin because multiple biopsies were obtained for those tissues. For ID 103, bars are below 100 to indicate viral DNA was below the limit of detection.

MPXV DNA was identified within MFI tissues in both cases in which infection progressed beyond 5 days post-inoculation. These MFI tissues included the placenta, decidua, and fetal membranes, and in the one animal where they were assayed, in the amniotic and chorionic membranes (Fig. 2). MPXV DNA was more widely disseminated within maternal tissues in the dams where infection progressed to 14 and 25 days post-inoculation, respectively, compared to the pregnancy where infection only progressed to 5 days post-inoculation (Fig. 2). The highest maternal tissue viral DNA load occurred in a biopsy of a skin lesion (Fig. 2). Placental tissues had infectious MPXV particles identified by plaque assay, with 2-9 × 10^6^ PFU/g of tissue identified from both pregnancies with fetal demise, but no infectious particles were detected in the amniotic fluid (Fig. S3, A and B). No fetal tissues were available for plaque assay due to very small fetal sizes. These data suggest that MPXV disseminates in maternal tissues by 5 days post-inoculation and reaches the MFI and fetus by 14-25 days post-inoculation.

### MPXV localizes to the placenta, fetal membranes, and chorionic plate

We identified locations where MPXV mRNA and antigens localize within the maternal-fetal interface by ISH and IHC, respectively. MPXV protein and mRNA were identified in the maternal-fetal interface (Fig. 3). While MPXV was not visually detectable in the fetal membranes from ID 101, there was intense focal localization in the fused fetal membranes from ID 102 (Fig. 3, A to D). Likewise, there was no localization of MPXV in the chorionic plate with ID 101 but there was scattered IHC localization of MPXV in ID 102 (Fig. 3, E to H). In ID 101, there was one distinct region of intense localization of MPXV mRNA and protein in the decidua basalis, but only a few scattered positive cells within the decidua otherwise (Fig. 3, I and K).

**Fig 3.**
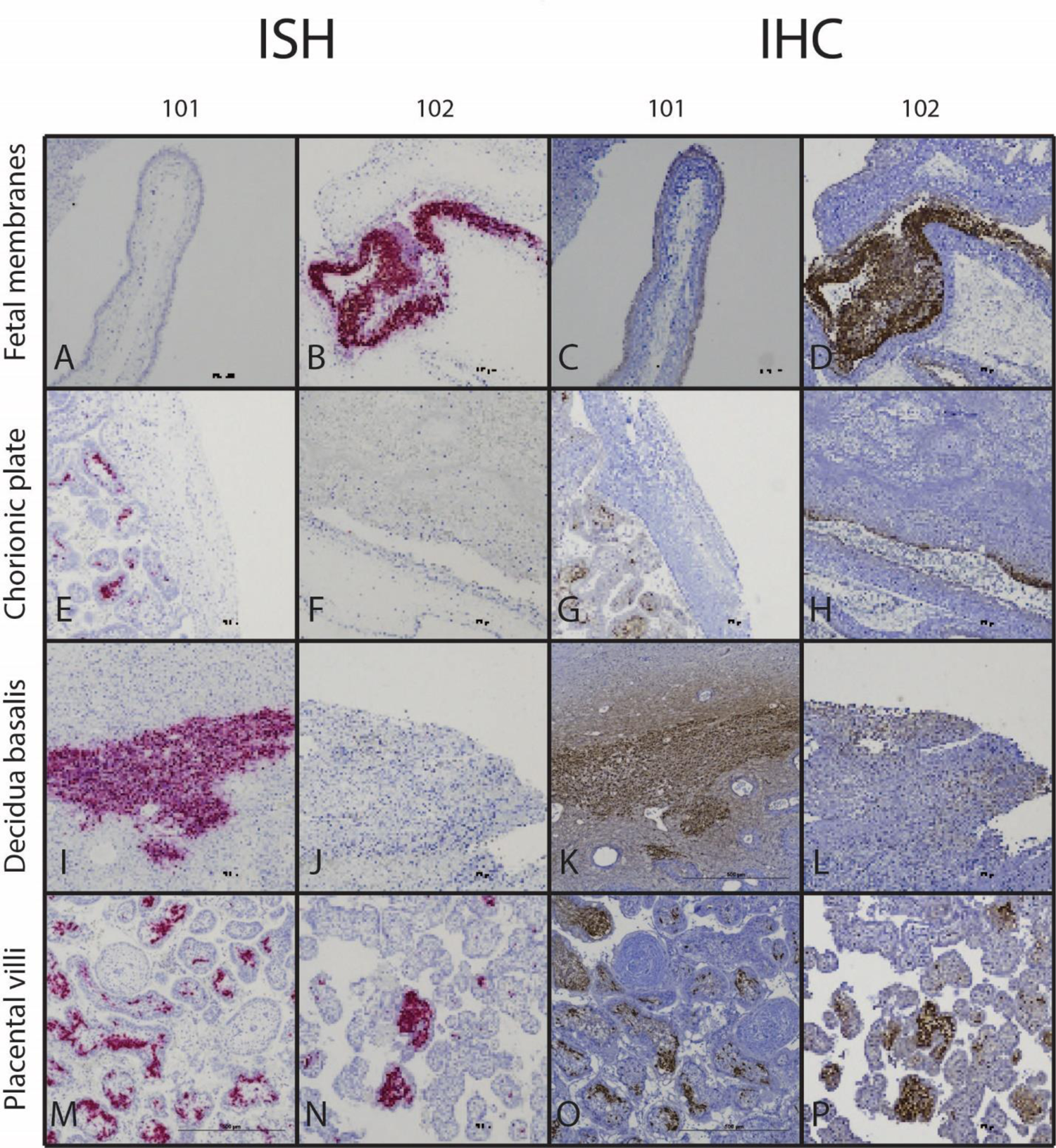
MPXV mRNA and antigen localization within maternal-fetal interface tissues by ISH and IHC. Representative images from ID 101 and 102 are presented. Fetal membranes (**A-D**), chorionic plate (**E-H**), decidua basalis (**I-L**), and placental villi (**M-P**) are shown. Images of the full tissue sections from which each of these photomicrographs were derived are shown in Figs. S4 and S5.

With ID 102, there was some localization of MPXV protein by IHC, however there was no detectable ISH signal in near sections of that specimen (Fig. 3, J and L). Finally, within the placental villi, there was intense localization of MPXV in the villous stroma but no apparent localization in the syncytiotrophoblasts or villous cytotrophoblasts in IDs 101 and 102 (Fig. 3, M to P, Figs. S4 and S5). We did not evaluate for MPXV antigen or RNA localization in the MFI of ID 103 because DNA viral loads were negative.

The intense localization of MPXV mRNA and protein in the stroma of the placental villi prompted further IHC to provide insight into the specific infected cells. Immunofluorescent staining for the Hofbauer cell (HBC) marker CD163 demonstrated the abundant presence of HBCs in the villous stroma, however only a minor subset of HBCs appeared to co-immunostain for MPXV proteins (Fig. 4A). Co-staining with anti-vimentin, which will stain both HBCs and villous mesenchymal fibroblasts, revealed more widespread co-staining, including of cells with distinctly fibroblastic morphology, suggesting that villous fibroblasts were also infected by MPXV (Fig. 4B). Finally, the villous mesenchyme includes placental blood vessels which are part of the fetal circulation. Co-staining with anti-CD31 identified a subset of endothelial cells that were also positive for MPXV protein (Fig. 4C). In rhesus macaques, the apical surface of the syncytiotrophoblasts also expresses CD31 (*15*). CD31 staining was seen on the apical surface of the villous syncytiotrophoblasts, however there was no co-localization of MPXV in the syncytium. There was also CD31 immunostaining in some places in the intervillous space, which likely reflects platelet-derived CD31 in intervillous fibrin.

**Fig 4.**
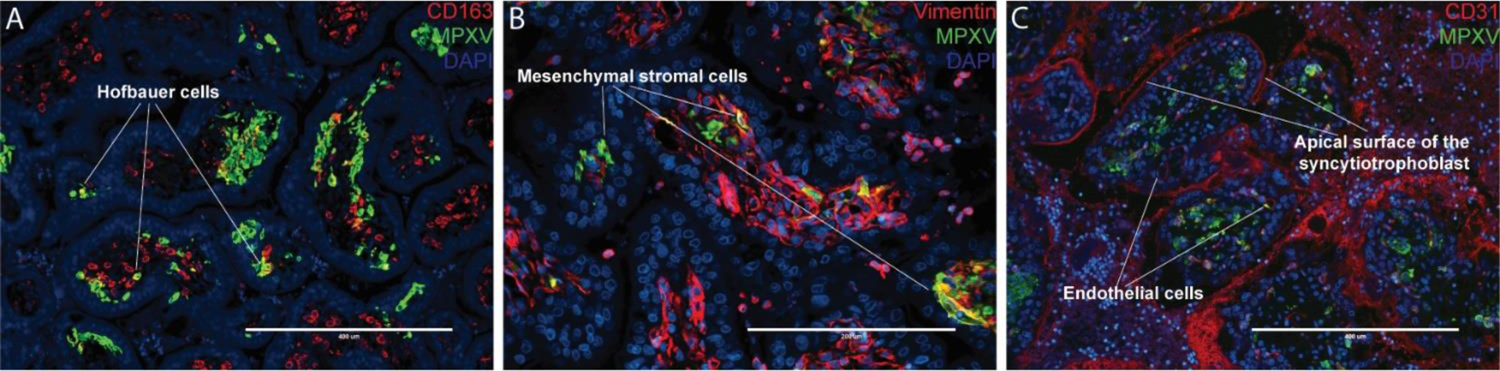
MPXV co-localizes with cell-specific markers in the placental villi. (**A**) MPXV co-localizes with CD163, a Hofbauer cell marker. MPXV is seen in few Hofbauer cells (CD163 staining) within the villi (not the majority of Hofbauer cells). (**B**) MPXV co-localization with vimentin, a marker for villous stromal cells. (**C**) MPXV co-localization with CD31, an endothelial cell marker which is also expressed on the apical surface of rhesus syncytiotrophoblasts. Representative placental images from ID 101 are shown.

The placentas of IDs 101 and 102 demonstrated few signs of tissue damage or histiocytic accumulation outside the areas where MPXV antigens were present. Notably, decidual spiral artery vasculitis was mild to absent with MPXV infection (Fig. 5A). Chorioamnionitis and chronic villitis were absent, but an increase in villous stromal macrophages were evident (Fig. 5B). Endovascular extravillous trophoblasts were negative for MPXV (Fig. 5C). There were no placental infarctions, but ID 101 did have a single focus of MPXV infection associated with decidual necrosis (Fig. 5, D and E). In summary, the amount of MPXV infection at the maternal-fetal interface was significant but does not appear to cause decidual vasculitis or lead to placental infarctions at this gestational age.

**Fig 5.**
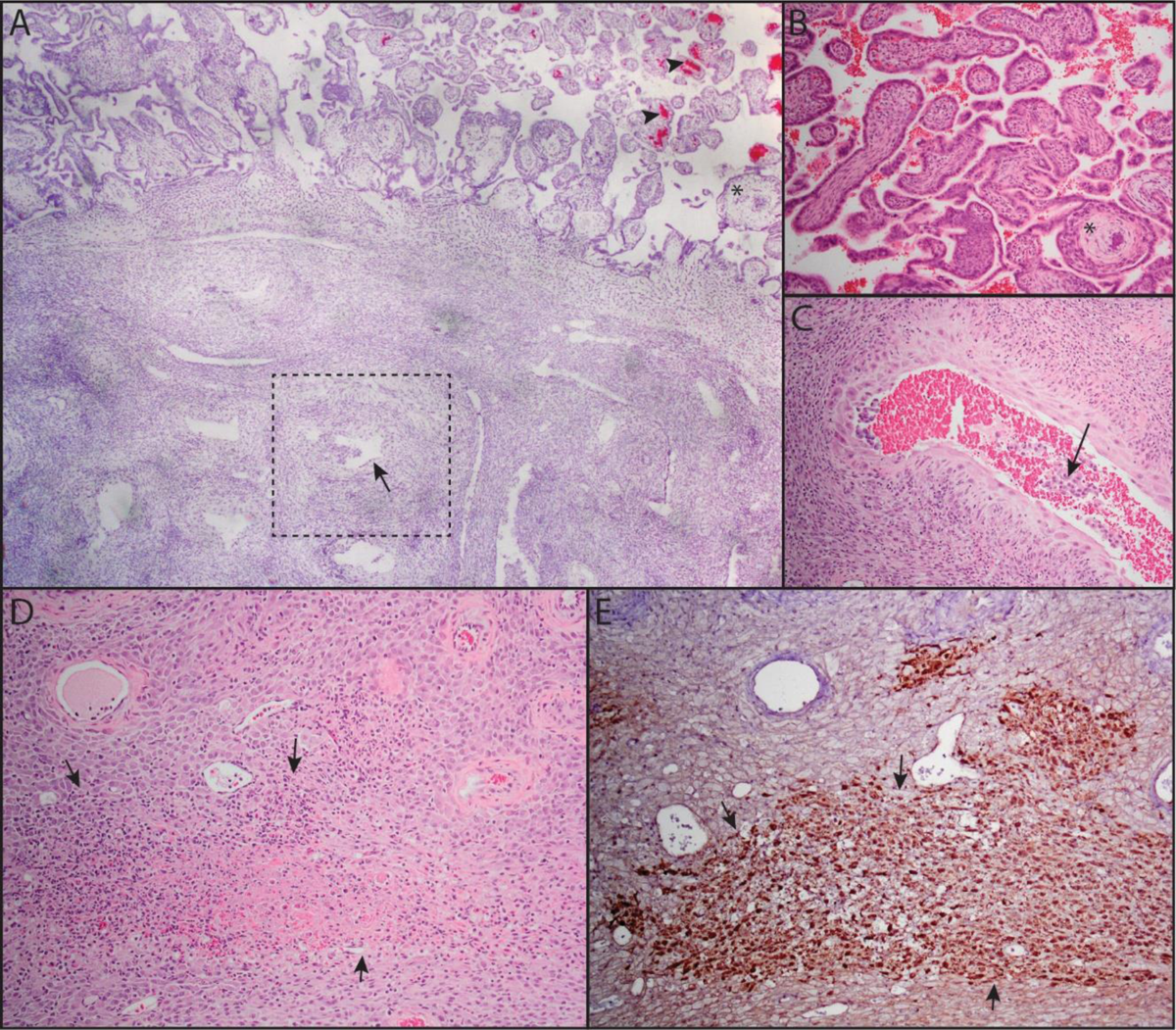
MPXV does not cause villitis or decidual vasculitis. (**A**) The placentas of IDs 101 and 102 demonstrated areas of villous stromal histiocytic accumulation associated with MPXV detection by ISH (red chromogen, arrow heads). The placental epithelium and endovascular extravillous trophoblasts invading the decidual spiral arteries (square) are negative for virus (arrow). There is no acute or chronic villitis (**B**) and the decidual spiral arteries are negative for leukocytoclastic vasculitis (arrow) (**C**). One of the animals (101) showed focal decidual necrosis (**D**) that co-labeled for MPXV virus (arrows outlining brown chromogen, **E**). * is a landmark in the upper right corner of panel A to compare image in A (2.5x objective) with image B (10x obj). Images C-E were all photographed with 10x objective.

### MPXV localizes to fetal tissues

We evaluated tissue localization of MPXV mRNA in fetal tissues by ISH to complement the determination of viral burden by qPCR. We identified positive ISH staining in multiple tissues from both cases of fetal demise (Fig. 6) with ISH staining in the neural tissue and skin of the head. Both fetuses had ISH staining in the lungs, liver, intercostal muscles, thoracic pleura, diaphragm, and the meninges and dura of the spinal cord. No significant histologic lesions were identified in any fetal tissues by qualitative histopathological analysis (table S1).

**Fig 6.**
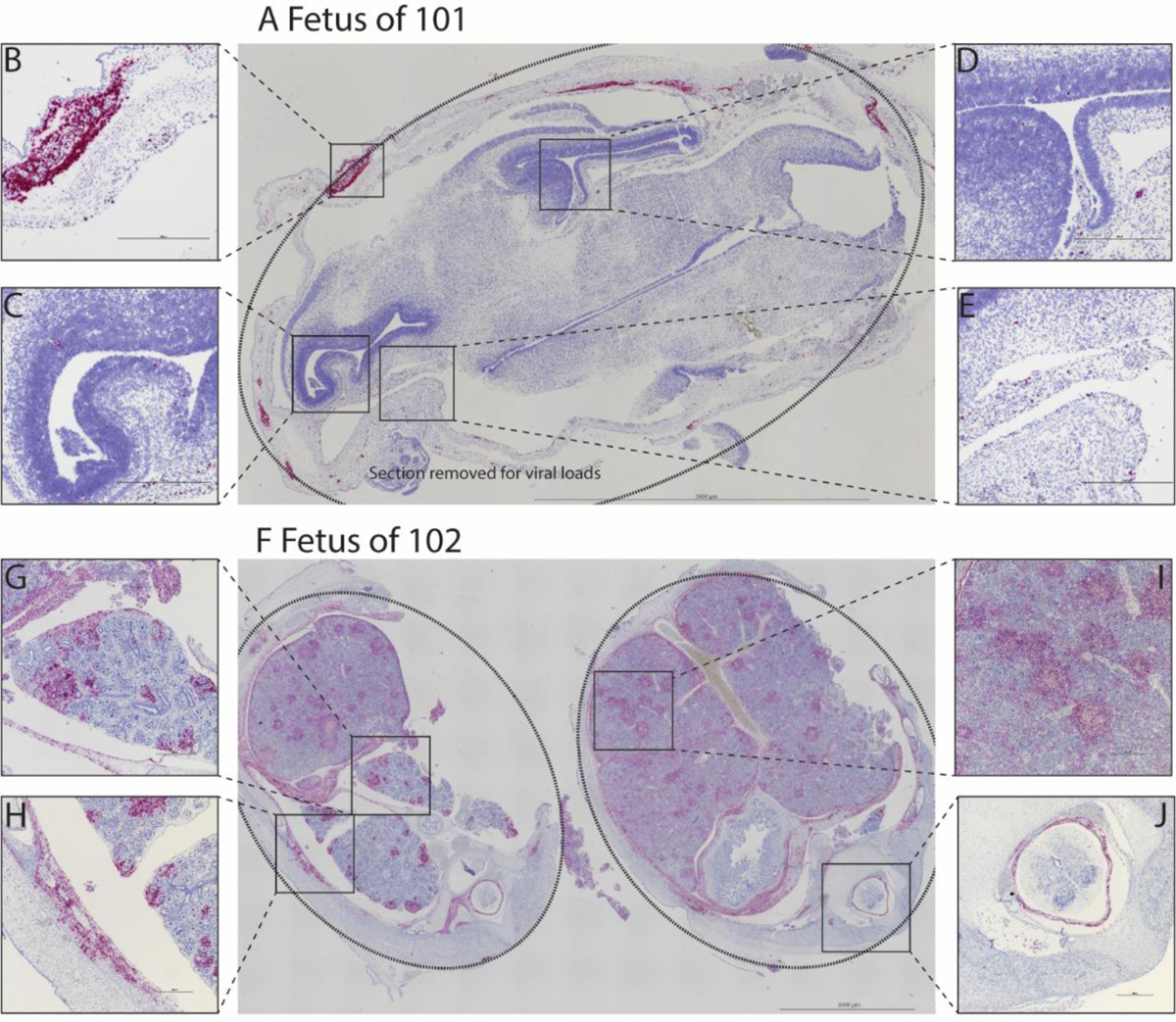
Fetal tissue MPXV ISH. Multiple regions within the head from the fetus of ID 101 (**A**) show ISH signal. The dotted line shows the head shape. Note: an area of neuropil was removed for viral load measurements. ISH signal was noted in the skin (**B**), in the subventricular zone of the anterior horn (**C**) and the posterior horn (**D**) of the lateral ventricle, and in the medial neuropil (**E**). Multiple regions within the torso from the fetus of ID 102 (**F**) show ISH signal. The dotted line shows the torso shape. ISH signal was found in the lung (**G**), intercostal muscles (**H**), liver (**I**), and in the meninges and dura surrounding the spinal cord (**J**).

## DISCUSSION

Our study establishes that clade IIb MPXV can be vertically transmitted in a translational macaque model of pregnancy, similar to clade I MPXV in human infection (*6,7*). This work also demonstrates that MPXV vertical transmission may occur after the appearance of skin lesions, which provides an opportunity for designing studies to determine the efficacy of antiviral medication and maternal vaccination to block vertical transmission. The pathway that the virus takes through the maternal-fetal interface to reach the fetus involves the decidua, the fetal membranes, and several cell populations within the stroma of the placental villi. MPXV is localized within multiple cell types in the placental villi, including Hofbauer cells, mesenchymal stromal cells, and endothelial cells. There was no decidual spiral artery vasculitis or MPXV antigens present in endovascular extravillous trophoblasts, unlike Zika virus infection, which does involve these important placental cells responsible for regulating decidual spiral artery remodeling and uteroplacental blood flow (*16*). Cases of fetal demise had MPXV DNA identified in multiple tissues, suggesting a hematogenous route of infection rather than selective permissiveness of a restricted population of fetal cells. Despite identifying MPXV DNA in multiple fetal tissues, there was an unexpected absence of significant histologic lesions and no inflammation within fetal tissues. This may be because the viral load rapidly reached high levels in the fetus causing demise without significant tissue damage. Further time course studies will resolve these questions. Understanding the pathogenesis of MPXV infection within the placenta and fetus that led to fetal demise in two of three pregnancies will help identify potential therapeutic targets for both clade I and clade IIb MPXV infections during pregnancy.

This macaque model of clade IIb MPXV infection is novel, in that it more closely models the natural history of human infection than other macaque models of clade IIb MPXV infection. For example, previous macaque studies used aerosol administration of lethal doses of MPXV which is not anticipated to be a physiological route of infection during this outbreak. We specifically selected a lower dose inoculum than previous macaque models (*17–20*) so that the dams could develop characteristic skin lesions without a high rate of mortality, which is what is observed in human infections (*1*). Determining whether vertical transmission occurs in an infection model that closely represents human infection is more informative for translational and therapeutic applications of this model. Additional clinical studies are needed to determine whether vertical transmission occurs in human clade IIb MPXV infection, and work with this translational model may be able to inform the design of human clinical studies that evaluate pregnancy outcomes and vertical transmission. Further work will be done in this rhesus macaque model to define the time course and pathway of MPXV to the fetal compartment and determine whether antiviral treatment blocks vertical transmission.

## MATERIALS AND METHODS

### Study design

All studies were conducted at the Wisconsin National Primate Research Center (WNPRC). Three pregnant Indian-origin rhesus macaques (Macaca mulatta) were inoculated intradermally between the scapulae with approximately 1.5 x10^5^ plaque forming units (PFU) of MPXV (hMPXV/USA/MA001/2022 (Lineage B.1, clade IIb)) in the first trimester. This dose was selected to mimic human skin-to-skin transmission while still ensuring infection in a rhesus macaque and was intermediate between the viral load present in a human skin lesion and a lethal intravenous dose in cynomolgus macaques (*21, 22*). Dams were free of Macacine herpesvirus 1 (Herpes B), simian retrovirus type D (SRV), simian T-lymphotropic virus type 1 (STLV), and simian immunodeficiency virus (SIV). Blood was collected for viral load testing and skin lesions were counted and swabbed for MPXV DNA according to the schedule in Table 1. The exact number of skin lesions was documented up to 100 lesions or indicated as >100 if greater than 100 lesions were present. Representative lesions were photographed. Fetal heart rate was monitored by doppler ultrasound during each sedation event. The dams were euthanized at the end of the study for collection of fetal, maternal-fetal interface, and maternal tissues.

**Table 1.** Timeline of dam inoculation, blood draw, skin lesion counts, fetal heart rate measurement, and skin lesion swab schedule. *Not done due to early study termination.

| Animal ID | Gestational day at inoculation | Blood draw, lesion count, fetal heart rate measurement frequency | Skin lesion swab schedule (days post-inoculation) |
| --- | --- | --- | --- |
| 101 | 31 | 2-3 times per week | 7, 9, 12, 14 |
| 102 | 32 |  | 7, 11, 14, 20, 25 |
| 103 | 34 |  | Not done* |

### MPXV stock

The mpox virus (MPXV) clade IIb stock used for this study was expanded from a vial of Monkeypox Virus, hMPXV/USA/MA001/2022 (Lineage B.1, clade IIb). The reagent was deposited by the Centers for Disease Control and Prevention and obtained from BEI Resources (item NR-58622). Briefly, Vero E6 cells (ATCC CRL-1586) were inoculated with stock virus at a multiplicity of infection (MOI) of 0.01. Before inoculation the virus stock was sonicated using a Q-Sonica cuphorn sonicator at 70% power for 30 seconds followed by a 15 second rest. This sonication was repeated for a total of 4x 45-second cycles. Following inoculation, cells were monitored daily. Approximately 80% cytopathic effect (CPE) was observed 48 hours later, at which time virus was harvested by collecting and pelleting cells. The pellet was resuspended in cold PBS and subsequently subjected to 3 freeze/thaw cycles and then sonicated for 4 cycles as described above to lyse cells and release virus. Virus was purified over a 36% sucrose cushion.

### MPXV stock sequencing and variant calling

Inactivated MPXV stock gDNA was processed for sequencing using the Oxford Nanopore Technologies SQK-RAD114 rapid sequencing kit. This kit randomly fragments the gDNA and adds adapters required for sequencing to the ends of the fragments. The stock gDNA concentration was 8 ng/ul, and a total of 80 ng of gDNA was processed for sequencing. The processed library was run on an R10.4.1 flow cell for 72 hours under default run conditions. 344,233 reads with a mean length of 2,888bp were obtained. Reads were imported into Geneious Prime® 2023.2.1 (Biomatters Ltd). The minimap2 mapping algorithm as implemented in this version of Geneious was used to map the reads to the expected MPXV reference sequence (GenBank accession: ON563414) with the following parameters: -x map-ont --frag=yes --eqx -m 60 --secondary=no. The -m override was important to avoid non-specific read mapping to the reference sequence. Within-sample variants were identified using the Geneious Find Variants/SNPs tool with the following key parameters: minimum coverage: 20; minimum variant frequency: 0.25; Maximum P-value: 10e-60.

### Ethics statement

The dams were cared for by the staff at the WNPRC in accordance with the regulations outlined in the US Animal Welfare Act, the principles described in the National Research Council’s Guide for the Care and Use of Laboratory Animals (*23*), and the recommendations of the Weatherall report (*24*). The University of Wisconsin - Madison approved this work under Institutional Biosafety Committee protocols B00000432, B00000764 and B00000182, and the Institutional Animal Care and Use Committee protocol G006670.

### Care and use of macaques

Female macaques were co-housed with a compatible male and observed daily for menses and breeding. Pregnancy was detected by abdominal ultrasound, and gestational age was estimated as previously described (25). For physical examinations, virus inoculations, ultrasound examinations, blood or swab collections, the dams were anesthetized with an intramuscular dose of ketamine (10 mg/kg) and monitored regularly until fully recovered from anesthesia. Blood samples from the femoral or saphenous vein were obtained using a vacutainer system or needle and syringe. The pregnant dams were individually housed in an enclosure with required floor space and fed using a nutritional plan based on recommendations published by the National Research Council (*24*). They were fed a fixed formula, extruded dry diet with adequate carbohydrate, energy, fat, fiber, mineral, protein, and vitamin content. The dry diet was supplemented with fruits, vegetables, and other edible enrichment (e.g., nuts, cereals, seed mixtures, yogurt, peanut butter, popcorn, etc.) to provide variety to the diet and to reinforce species-specific behaviors such as foraging. To further promote psychological well-being, they were provided with food enrichment, structural enrichment, and/or manipulanda. The dams were evaluated by trained animal care staff at least twice each day for signs of pain, distress, or illness by observing appetite, stool quality, activity level, and physical condition. Any abnormal presentation prompted examination by a veterinarian.

### Whole blood preparation

Whole blood was collected in EDTA-treated vacutainer tubes. The EDTA tubes were inverted and whole blood removed, aliquoted, and stored at −80°C until DNA qPCR.

### Viral DNA (vDNA) isolation and quantitative polymerase chain reaction (qPCR)

Viral DNA was extracted from 300 ul fluid samples (including whole blood, urine, CSF, amniotic fluid and eluates from swab samples) using the Viral Total Nucleic Acid Purification Kit for the Maxwell RSC instrument (Promega, Madison WI) with the following modifications. After the addition of the lysis solution, samples were incubated for 10 minutes at room temperature and then at 80°C for 10 minutes.

Viral DNA was quantified using the E9L_NVAR qPCR assay targeting the DNA polymerase gene of non-variola orthopox viruses (*26*) with the Taqman Fast Advanced Master Mix (LifeTechnologies, Carlsbad, CA). DNA quantification was accomplished by interpolation onto a standard curve consisting of ten-fold serial dilutions of the RGTM 10223 plasmid provided by National Institute of Standards and Technology that contains the target region of the assay. The qPCR reaction contains 500 nM of each forward (5’-TCAACTGAAAAGGCCATCTATGA-3’) and reverse primer (5’-GAGTATAGAGCACTATTTCTAAATCCCA-3’) and 200 nM probe (5’-CCATGCAATATACGTACAAGATAGTAGCCAAC-3’). The cycling conditions are as follows: 95°C for 5 min followed by 50 cycles of 95°C for 15 seconds, 60°C for 1 minute and 72°C 1 second. The lower limit of detection for this assay is 700 DNA copy Eq/ml. DNA was extracted from up to 100 mg tissue biopsies. Samples were weighed prior to homogenization in 1 ml Trizol (Invitrogen, Waltham, MA) using the Qiagen TissueLyser II.

DNA was recovered from the interphase layer following phase separation using bromochloropropane and was ethanol precipitated, washed with 0.1 M sodium citrate in 10% ethanol. DNA was then pelleted and re-suspended in 8mM sodium hydroxide, buffered with HEPES to ∼ pH 7.2. Viral DNA was then quantified by qPCR as described above.

### Viral quantification by plaque assay

Assays for replication competent MPXV quantification were performed via plaque assay on confluent monolayers of Vero E6 cells (ATCC CRL-1586). The presence of replication competent virus was assessed in eluates from skin lesion swabs, amniotic fluid collected at necropsy, and homogenates of placental tissues collected at necropsy. Prior to beginning plaque assays, fluid samples (skin lesion swab eluates and amniotic fluid) were either subjected to 3 rounds of sonication using a Fisherbrand Model 505 sonicator with a cup-horn attachment at 70% power for 30 seconds with a 15 second rest in between each round (ID 101) or were not sonicated (ID 102). We assessed the impact of sonication on viral titer in a lesion swab sample from ID 101 and determined sonication had a negligible impact on viral titer (Fig S3C). Prior to plaque assay, placental tissues were weighed, placed in 1mL of infection media (1X DMEM (Gibco)-2% fetal bovine serum) and homogenized using a Dounce tissue grinder to generate a tissue slurry. This tissue slurry was then sonicated using the same parameters described above and clarified by centrifugation at 400xg for 10 seconds. Each 0.1mL aliquot of fluid or tissue slurry supernatant was serially diluted 10-fold in infection media and inoculated on to Vero E6 cells in duplicate in 12-well culture plates. The plates were incubated for 1 hour at 37°C 5% CO_2_ with gentle rocking every 15 minutes to allow for virus adsorption and to ensure even distribution of sample across the monolayer. Following incubation, the monolayers were overlaid with 1mL of overlay media containing a 1:1 mixture of 2.4% microcrystalline cellulose (Beantown Chemical, Hudson, NH, USA) and 2X DMEM (Gibco) with 0.5% bovine serum albumin (v/v), 5% 1M HEPES, 2% GlutaMax (Gibco), and 1% penicillin/streptomycin/amphotericin B solution (Gibco). The plates were incubated for 72 hours at 37°C 5% CO_2_ to allow for plaque formation. After 72 hours, the overlay media was discarded, and the cells were fixed with ice-cold ethanol for 20 minutes at room temperature. The monolayers were stained with 0.5% crystal violet for 10 minutes at room temperature and the number of plaques was counted.

### RNA in-situ hybridization (ISH)

To detect Mpox messenger RNA (mRNA) in formalin-fixed paraffin-embedded (FFPE) tissues, ISH was performed as described previously using the RNAscope® 2.5 HD Detection Kit (RED) for FFPE Tissues (Advanced Cell Diagnostics, Newark, CA, USA) (*27*). Briefly, an ISH probe targeting fragment 2-1292 of MPXV-specific gene *D1L*, which aligns to 188309-189622 of KJ642618.1, was designed and synthesized by Advanced Cell Diagnostics (Cat#534671). Tissue sections were deparaffinized with xylene, underwent a series of ethanol washes and peroxidase blocking, and were then heated in kit-provided antigen retrieval buffer and digested by kit-provided proteinase. Sections were exposed to ISH probes and incubated at 40°C in a hybridization oven for 2 hours. After rinsing, ISH signal was amplified using kit-provided Pre-amplifiers and Amplifier conjugated to alkaline phosphatase, and then incubated with a Fast Red substrate solution for 10 minutes at room temperature. Sections were then counterstained with hematoxylin, air-dried, and coverslipped. Photomicrographs were taken on a Nikon Eclipse microscope.

### Immunohistochemistry (IHC)

Formalin-fixed, paraffin-embedded sections were subjected to a heat-induced epitope retrieval protocol in Tris-EDTA pH 9 buffer (Abcam, Waltham, MA, USA) at 110°C for 15 minutes.

Slides were blocked with 3% H_2_O_2_ for 10 minutes and with Background Punisher (Biocare Medical, Concord, CA, USA) for 30 minutes. Sections were immunohistochemically stained using an anti-vaccinia virus primary antibody at 1:1000 (Genetex, Irvine, CA, USA) (table S2) diluted in a 5% normal goat serum in Tris buffered saline with 1% Tween-20 and incubated for 30 minutes. An anti-rabbit polymer-horseradish peroxidase conjugated secondary antibody (Biocare Medical, Concord, CA, USA) was used undiluted and incubated 25 minutes at room temperature to detect bound antibody. Signal was visualized with Betazoid DAB Chromagen Kit (Biocare Medical, Concord, CA, USA) and counterstained with hematoxylin. All washes and reagent dilutions were in Tris-buffered saline pH 8.4 with 0.01% Tween-20 (Fisher BioReagents, Waltham, MA, USA). All incubations and washes were conducted at room temperature.

### Immunofluorescence (IF) microscopy

Tissue section slides were prepared from paraformaldehyde-fixed, paraffin-embedded blocks. Slides were deparaffinized in xylene and hydrated through an ethanol series. Antigen retrieval was performed in a Tris EDTA pH 9 buffer (Abcam, Waltham, MA, USA) via microwave for 8 min at ∼95°C. Slides were then blocked in a blocking buffer (2% goat serum, 1% bovine serum albumin, 0.1% Triton X-100 and 0.05% Tween-20 in Tris buffered saline) at room temperature for one hour. Primary antibodies (table S2) were diluted in blocking buffer and incubated on the slides overnight at 4°C. CD163 was used as a marker for Hofbauer cells, CD31 for endothelial cells and the apical surface of the syncytiotrophoblast (*15*), and vimentin for mesenchymal stromal cells. The following day, slides were incubated with secondary antibodies (table S2) for 1 hour at room temperature. Slides were coverslipped with a DAPI inclusive mounting media (Prolong Anti Fade, Thermofisher, Waltham MA, USA). Stained tissue sections were imaged using an EVOS AutoFL microscope system (Life Technologies, Grand Island, NY, USA). The light cubes used for fluorescent imaging were Texas Red, Cy5, and DAPI (Life Technologies, Grand Island, NY, USA) for detecting ALEXA 594, ALEXA 647, and DAPI, respectively. Control immunostaining was done with isotype controls (table S2) and imaging of the control immunostaining can be found in fig S6.

## List of Supplementary Materials

Fig S1 to S6

Tables S1 to S2

## Acknowledgments

We thank the Wisconsin National Primate Research Center (WNPRC) Veterinary Services, Scientific Protocol Implementation, Pathology Services, and Behavioral Services Unit staff for assistance with animal procedures, including breeding, monitoring, surgery, and necropsy. We thank the WNPRC Virology Services and Assay Services units for performing viral DNA isolation, MPXV DNA qRT-PCR, and the hormone assays. Dr. Leticia Reyes (UW-Madison Dept. of Pathobiological Sciences) provided assistance and guidance in EVOS immunofluorescence microscopy. We thank the University of Wisconsin Translational Research Initiatives in Pathology (TRIP) Laboratory, supported by the UW Department of Pathology and Laboratory Medicine, UWCCC (P30 CA014520) and the Office of Research Infrastructure Programs-NIH (S10 OD023526) for use of its facilities and services.

## Funding

National Institutes of Health grant R01AI182082 (ELM), National Institutes of Health grant P01AI132132 (DHO), National Institutes of Health grant 2R24OD017850 (DHO).

## Author contributions

Conceptualization: NPK, AMM, SB, ERR, TGG, ELM

Methodology: NPK, AMM, SB, SVC, HAS, AMW, TGG, ELM

Investigation: NPK, AMM, SB, XZ, NB, AM, AK, JAK, GD, GV, TM, SVC, HAS, PB, AMW, TGG, ELM

Visualization: NPK, AMM, TM, HAS, TGG, ELM

Funding acquisition: DHO, TGG, ELM

Project administration: ERR, TGG, ELM

Supervision: SVC, HAS, PB, AMW, DHO, TCF, TGG, ELM

Writing-original draft: NPK, AMM, SB, JAK, HAS, AMW, TGG, ELM

Writing-review and editing: NPK, AMM, SB, ERR, XZ, NB, AM, AK, JAK, GD, GV, TM, SVC, HAS, PB, AMW, DHO, TGG, ELM

## Competing interests

Authors declare that they have no competing interests.

## Data and materials availability

All raw data and code used for data analysis and figure making in this manuscript can be found at: https://go.wisc.edu/xw4o53

## Notes

### Competing Interest Statement

The authors have declared no competing interest.

## References

1. Mpox ongoing 2022 global outbreak cases and data (2024) (available at https://www.cdc.gov/poxvirus/mpox/response/2022/index.html).

2. D. A. Schwartz, S. Ha, P. Dashraath, D. Baud, P. R. Pittman, K. A. Waldorf, Monkeypox Virus in Pregnancy, the Placenta and Newborn: An Emerging Poxvirus with Similarities to Smallpox and Other Orthopoxvirus Agents Causing Maternal and Fetal Disease. Arch. Pathol. Lab. Med. (2023), doi:10.5858/arpa.2022-0520-SA.

3. J. G. Breman, Kalisa-Ruti, M. V., Steniowski, E., Zanotto, A. I., Gromyko, I., Arita, Human monkeypox, 1970-79. Bull. World Health Organ. 58, 165–182 (1980).

4. A. M. McCollum, I. K. Damon, Human monkeypox. Clin. Infect. Dis. 58, 260–267 (2014).

5. D. D. Djuicy, S. A. Sadeuh-Mba, C. N. Bilounga, M. G. Yonga, J. B. Tchatchueng-Mbougua, G. D. Essima, L. Esso, I. M. E. Nguidjol, S. F. Metomb, C. Chebo, S. M. Agwe, P. A. Ankone, F. N. N. Ngonla, H. M. Mossi, A. G. M. Etoundi, S. I. Eyangoh, M. Kazanji, R. Njouom, Concurrent Clade I and Clade II Monkeypox Virus Circulation, Cameroon, 1979-2022. Emerg. Infect. Dis. 30, 432–443 (2024).

6. P. K. Mbala, J. W. Huggins, T. Riu-Rovira, S. M. Ahuka, P. Mulembakani, A. W. Rimoin, J. W. Martin, J.-J. T. Muyembe, Maternal and Fetal Outcomes Among Pregnant Women With Human Monkeypox Infection in the Democratic Republic of Congo. J. Infect. Dis. 216, 824–828 (2017).

7. P. R. Pittman, J. W. Martin, P. M. Kingebeni, J.-J. M. Tamfum, G. Mwema, Q. Wan, P. Ewala, J. Alonga, G. Bilulu, M. G. Reynolds, X. Quinn, S. Norris, M. B. Townsend, P. S. Satheshkumar, J. Wadding, B. Soltis, A. Honko, F. B. Güereña, L. Korman, K. Patterson, D. A. Schwartz, J. W. Huggins, Kole Human Mpox Infection Study Group, Clinical characterization and placental pathology of mpox infection in hospitalized patients in the Democratic Republic of the Congo. PLoS Negl. Trop. Dis. 17, e0010384 (2023).

8. C. Besombes, F. Mbrenga, C. Malaka, E. Gonofio, L. Schaeffer, X. Konamna, B. Selekon, J. Namsenei-Dankpea, C. Gildas Lemon, J. Landier, C. von Platen, A. Gessain, J. C. Manuguerra, A. Fontanet, E. Nakouné, Investigation of a mpox outbreak in Central African Republic, 2021-2022. One Health 16, 100523 (2023).

9. CDC, 2023 outbreak in Democratic Republic of the CongoCenters for Disease Control and Prevention (2024) (available at https://www.cdc.gov/poxvirus/mpox/outbreak/2023-drc.html).

10. L. P. Oakley, K. Hufstetler, J. O’Shea, J. D. Sharpe, C. McArdle, V. Neelam, N. M. Roth, E. O. Olsen, M. Wolf, L. Z. Pao, J. A. W. Gold, K. M. Davis, D. Perella, S. Epstein, M. K. Lash, O. Samson, J. Pavlick, A. Feldpausch, J. Wallace, A. Nambiar, V. Ngo, U.-A. Halai, C. W. Richardson, T. Fowler, B. P. Taylor, J. Chou, L. Brandon, R. Devasia, E. K. Ricketts, C. Stockdale, M. Roskosky, R. Ostadkar, Y. Vang, R. R. Galang, K. Perkins, M. Taylor, M. J. Choi, P. J. Weidle, P. Dawson, S. Ellington, CDC Mpox Analytics Team, Mpox Cases Among Cisgender Women and Pregnant Persons - United States, May 11-November 7, 2022. MMWR Morb. Mortal. Wkly. Rep. 72, 9–14 (2023).

11. M. M. Sampson, G. Magee, E. A. Schrader, K. L. Dantuluri, A. Bukhari, C. Passaretti, L. Temming, M. Leonard, J. B. Philips, D. Weinrib, Mpox (Monkeypox) Infection During Pregnancy. Obstet. Gynecol. 141, 1007–1010 (2023).

12. A. Khalil, A. Samara, P. O’Brien, C. M. Coutinho, G. Duarte, S. M. Quintana, S. N. Ladhani, Monkeypox in pregnancy: update on current outbreak. Lancet Infect. Dis. 22, 1534–1535 (2022).

13. M. Vouga, K. Nielsen-Saines, P. Dashraath, D. Baud, The monkeypox outbreak: risks to children and pregnant women. Lancet Child Adolesc Health (2022), doi:10.1016/S2352-4642(22)00223-1.

14. M. L. Fahrni, Priyanka, O. P. Choudhary, Possibility of vertical transmission of the human monkeypox virus. Int. J. Surg. 105, 106832 (2022).

15. G. I. Bondarenko, D. W. Burleigh, M. Durning, E. E. Breburda, R. L. Grendell, T. G. Golos, Passive immunization against the MHC class I molecule Mamu-AG disrupts rhesus placental development and endometrial responses. J. Immunol. 179, 8042–8050 (2007).

16. M. R. Koenig, A. M. Mitzey, X. Zeng, L. Reyes, H. A. Simmons, T. K. Morgan, E. K. Bohm, J. C. Pritchard, J. A. Schmidt, E. Ren, F. B. Leyva Jaimes, E. Winston, P. Basu, A. M. Weiler, T. C. Friedrich, M. T. Aliota, E. L. Mohr, T. G. Golos, Vertical transmission of African-lineage Zika virus through the fetal membranes in a rhesus macaque (Macaca mulatta) model. PLoS Pathog. 19, e1011274 (2023).

17. A. Zuiani, C. L. Dulberger, N. S. De Silva, M. Marquette, Y.-J. Lu, G. M. Palowitch, A. Dokic, R. Sanchez-Velazquez, K. Schlatterer, S. Sarkar, S. Kar, B. Chawla, A. Galeev, C. Lindemann, D. A. Rothenberg, H. Diao, A. C. Walls, T. A. Addona, F. Mensa, A. B. Vogel, L. M. Stuart, R. van der Most, J. R. Srouji, Ö. Türeci, R. B. Gaynor, U. Şahin, A. Poran, A multivalent mRNA monkeypox virus vaccine (BNT166) protects mice and macaques from orthopoxvirus disease. Cell 187, 1363–1373.e12 (2024).

18. A. T. Russo, A. Berhanu, C. B. Bigger, J. Prigge, P. M. Silvera, D. W. Grosenbach, D. Hruby, Co-administration of tecovirimat and ACAM2000TM in non-human primates: Effect of tecovirimat treatment on ACAM2000 immunogenicity and efficacy versus lethal monkeypox virus challenge. Vaccine 38, 644–654 (2020).

19. E. M. Mucker, J. D. Shamblin, A. J. Goff, T. M. Bell, C. Reed, N. A. Twenhafel, J. Chapman, M. Mattix, D. Alves, R. F. Garry, L. E. Hensley, Evaluation of Virulence in Cynomolgus Macaques Using a Virus Preparation Enriched for the Extracellular Form of Monkeypox Virus. Viruses 14 (2022), doi:10.3390/v14091993.

20. M. Saijo, Y. Ami, Y. Suzaki, N. Nagata, N. Iwata, H. Hasegawa, I. Iizuka, T. Shiota, K. Sakai, M. Ogata, S. Fukushi, T. Mizutani, T. Sata, T. Kurata, I. Kurane, S. Morikawa, Virulence and pathophysiology of the Congo Basin and West African strains of monkeypox virus in non-human primates. J. Gen. Virol. 90, 2266–2271 (2009).

21. A. Berhanu, J. T. Prigge, P. M. Silvera, K. M. Honeychurch, D. E. Hruby, D. W. Grosenbach, Treatment with the smallpox antiviral tecovirimat (ST-246) alone or in combination with ACAM2000 vaccination is effective as a postsymptomatic therapy for monkeypox virus infection. Antimicrob. Agents Chemother. 59, 4296–4300 (2015).

22. B. Huang, H. Zhao, J. Song, L. Zhao, Y. Deng, W. Wang, R. Lu, W. Wang, J. Ren, F. Ye, H. Tian, G. Wu, H. Ling, W. Tan, Isolation and Characterization of Monkeypox Virus from the First Case of Monkeypox - Chongqing Municipality, China, 2022. China CDC Wkly 4, 1019– 1024 (2022).

23. National Research Council, Division on Earth and Life Studies, Institute for Laboratory Animal Research, Committee on Guidelines for the Use of Animals in Neuroscience and Behavioral Research, Guidelines for the Care and Use of Mammals in Neuroscience and Behavioral Research (National Academies Press, 2003).

24. The Weatherall report on the use of non-human primates in research (available at https://royalsociety.org/topics-policy/publications/2006/weatherall-report).

25. E. L. Mohr, L. N. Block, C. M. Newman, L. M. Stewart, M. Koenig, M. Semler, M. E. Breitbach, L. B. C. Teixeira, X. Zeng, A. M. Weiler, G. L. Barry, T. H. Thoong, G. J. Wiepz, D. M. Dudley, H. A. Simmons, A. Mejia, T. K. Morgan, M. S. Salamat, S. Kohn, K. M. Antony, M. T. Aliota, M. S. Mohns, J. M. Hayes, N. Schultz-Darken, M. L. Schotzko, E. Peterson, S. Capuano 3rd, J. E. Osorio, S. L. O’Connor, T. C. Friedrich, D. H. O’Connor, T. G. Golos, Ocular and uteroplacental pathology in a macaque pregnancy with congenital Zika virus infection. PLoS One 13, e0190617 (2018).

26. Y. Li, V. A. Olson, T. Laue, M. T. Laker, I. K. Damon, Detection of monkeypox virus with real-time PCR assays. J. Clin. Virol. 36, 194–203 (2006).

27. J. Liu, E. M. Mucker, J. L. Chapman, A. M. Babka, J. M. Gordon, A. V. Bryan, J. L. W. Raymond, T. M. Bell, P. R. Facemire, A. J. Goff, A. Nalca, X. Zeng, Retrospective detection of monkeypox virus in the testes of nonhuman primate survivors. Nat Microbiol 7, 1980–1986 (2022).

