## Supplementary material for "Mpox virus (MPXV) vertical transmission and fetal demise in a pregnant rhesus macaque model": Fig S1-S6 and Table S1-S2

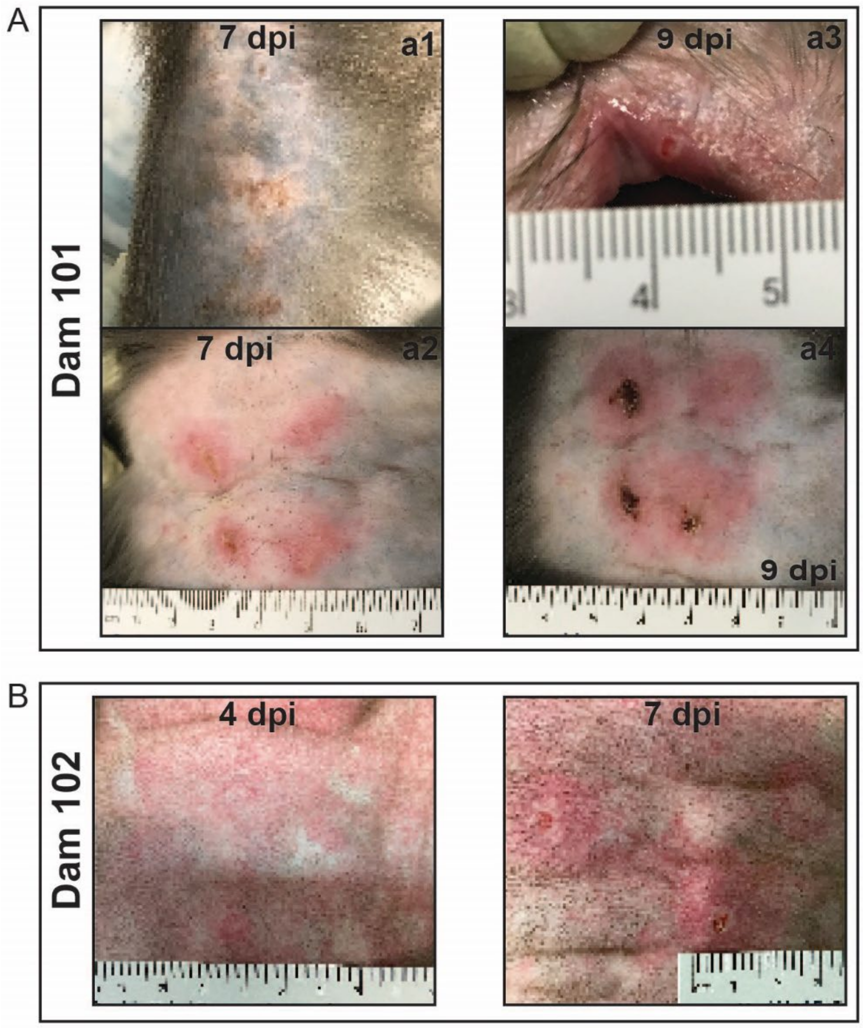


**Fig S1. Skin lesion appearance.** (**A**) Skin lesions first appeared on dam 101 at 7 days post-inoculation (dpi). (**a1**) Multiple pustules with superficial serocellular crusting appeared on an arm distant from the inoculation sites. (**a2**) Large pustules and dermal ulcerations surrounded by erythema appeared at the sites of inoculation between the scapula with smaller macules, papules, and pustules developing adjacent and between inoculation sites. (**a3**) By 9 dpi, there was focal ulceration of the oral mucocutaneous junction. (**a4**) By 9 dpi, injection site ulcers had developed eschars and were still surrounded by intense erythema as well as small pustules and ulcers. (**B**) Skin lesions first appeared on dam 102 at 4 dpi. The skin inoculation sites between the scapulae were erythematous macules and papules. By 7 dpi, the inoculation sites progressed to ulcers and a papule surrounded by erythema.

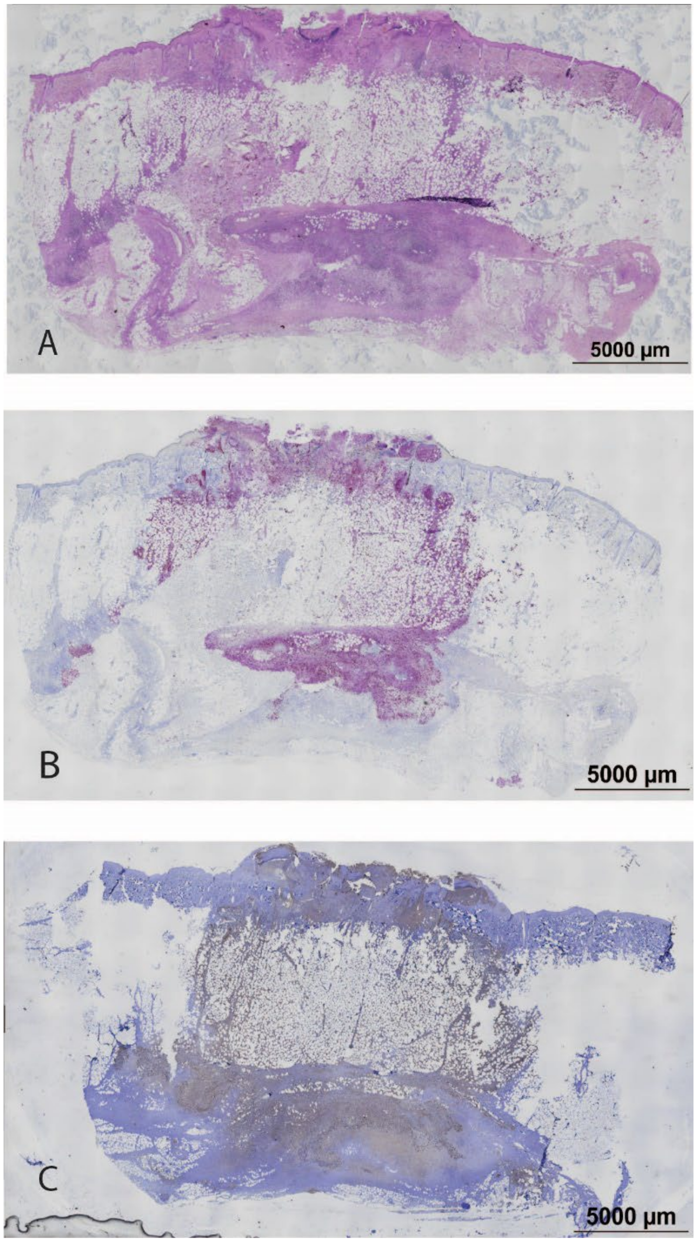


**Fig S2. Maternal skin lesion histology and MPXV loacalization by ISH and IHC.** (**A**) Hematoxylin and eosin staining of 5µM section. There is focally extensive epidermal and dermal ulceration, necrotizing dermatitis, steatitis of the underlying subcutaneous adipose, and severe diffuse neutrophilic, lymphoplasmyctic and occasionally eosinophilic and histiocytic panniculitis with multifocal vasculitis and vascular necrosis. The intact skin on either side of the ulcer has multiple epidermal pustules, ballooning degeneration of keratinocytes, and moderate to severe multifocal dermatitis. (**B**) ISH signal (red) and hematoxylin (blue) staining within the ulcer, dermis, subcutis, and panniculus of a serial skin section. (**C**) IHC signal (brown) and hematoxylin (blue) staining within the ulcer, dermis, subcutis, and panniculus of a nearby skin section.

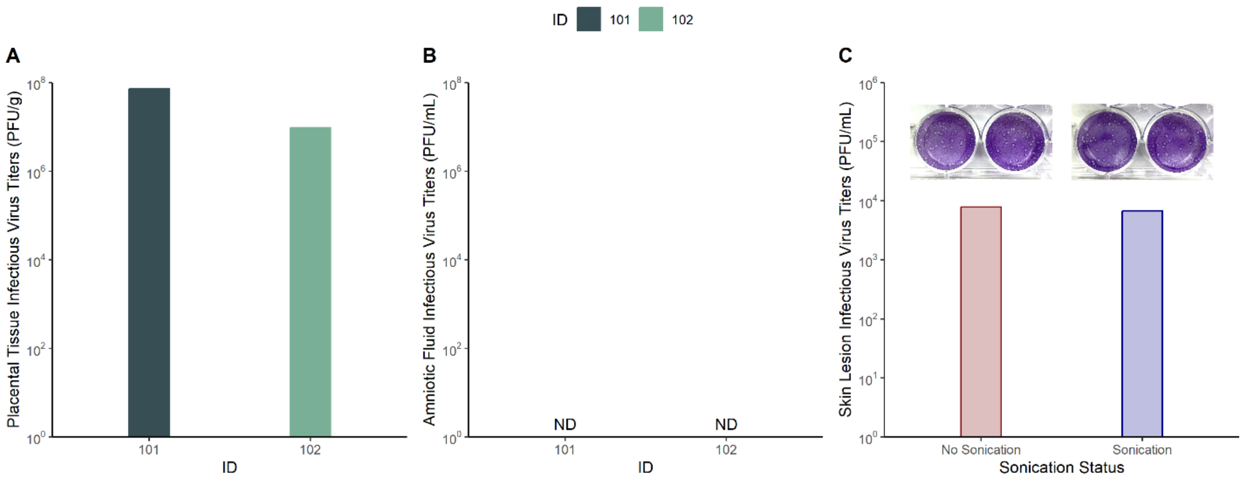


**Fig S3. Placental tissue and amniotic fluid infectious virus titers and sonication comparison.** MPXV infectious virus titers were measured via plaque assay at the time of necropsy in placental tissue homogenates (**A**) and amniotic fluid (**B**) for IDs 101 and 102. For ID 103, no plaque assays were performed because viral DNA titers were below the limit of detection. The impact of performing sonication on infectious virus titers was determined using an eluate from a skin lesion swab from ID 101 at 7 days post-inoculation (C). Representative images show plaque formation following a 1:10 dilution of sample that was either not sonicated or sonicated.

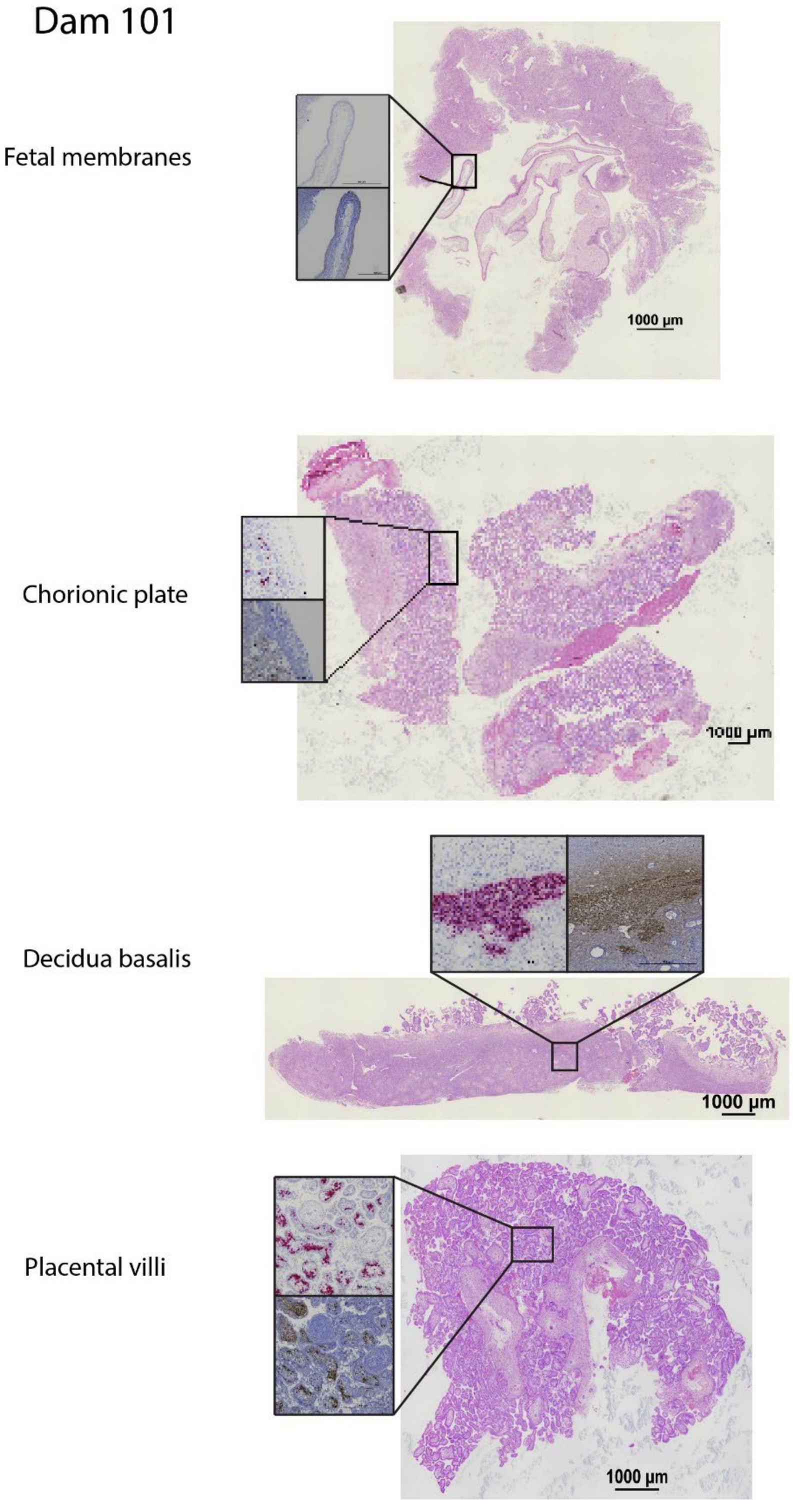


**Fig S4. Full thickness sections shown in figure 3 for dam 101 demonstrating the location where each insert was located in the full thickness section.**

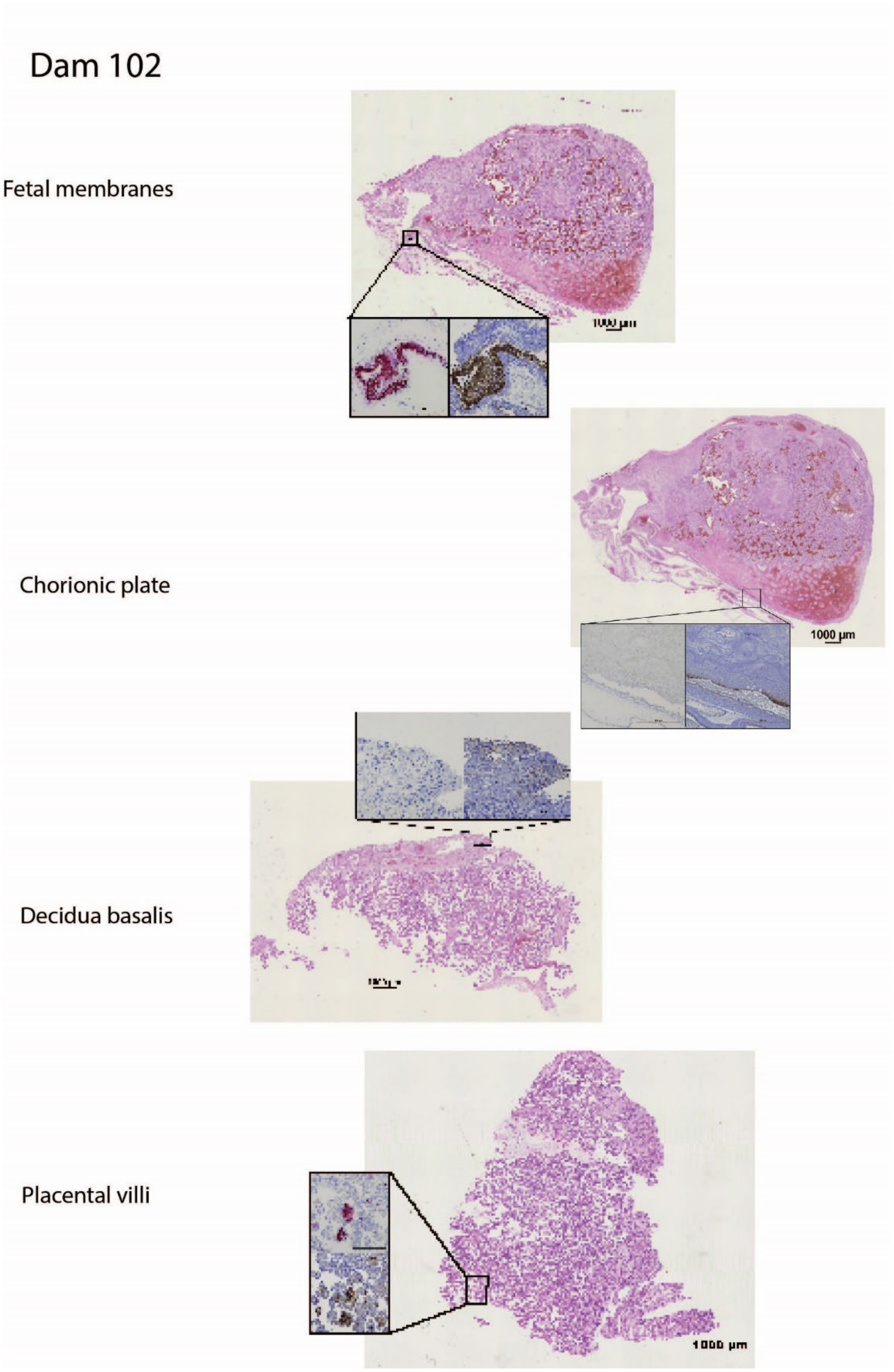


**Fig S5. Full thickness sections shown in figure 3 for dam 102 demonstrating the location where each insert was located in the full thickness section.**

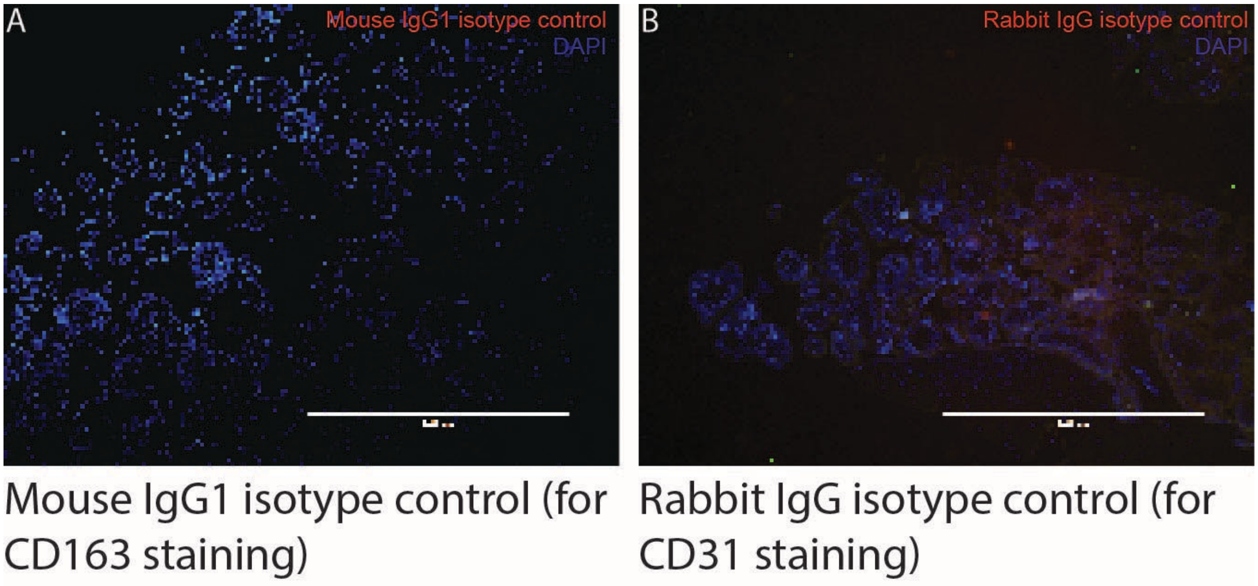


**Fig S6. Isotype control staining for immunofluorescence imaging.** (**A**) Placental tissue section stained with DAPI and mouse IgG1 antibody as an isotype control for the CD163 mouse IgG1 antibody. (**B**) Placental tissue section stained with DAPI and rabbit IgG antibody as an isotype control for the CD31 rabbit IgG antibody.

| | Tissue name(s) | Dam 101 & fetus | Dam 102 & fetus | Dam 103 and fetus |
|---|---|---|---|---|
| **Maternal** | Final summary comments | Dermal changes in this animal are consitent with pox viral infection as are the changes in the epidermis of the tongue. Mpox has been documented to replicate within lymph nodes consistent with the findings in the axillary, inguinal, and mesenteric lymph nodes.<br><br>Cardiac and pulmonary changes likely pre-date and are unrelated to the experimental manipulation in this animal. The cardiac changes are consistent with macaque hypertrophic cardiomyopathy which is often associated with acute unanticipated demise. The presence of multinucleate giant cells within the lungs is concerning. Special stains were evaluated and there was no evidence of acid-fast or fungal organisms. | Dermal changes in this animal are consistent with pox viral infection.<br><br>The cardiac changes are consistent with macaque hypertrophic cardiomyopathy which is often associated with acute unanticipated demise.<br><br>The histologic changes in the decidua parietalis of the fetal membranes and the placenta are significant. | Dermal changes in this animal are consistent with pox viral infection.<br><br>This animal has marked multifocal acute and subacute retroplacental hemorrhage, as well as severe hemorrhage within the conceptus. The cause of the impending fetal loss in this case is likely multifactorial with the mild neutrophilic endometritis and myometritis contributing to the MFI decompensation.<br><br>This animal has incidental chronic renal and cardiac changes. |
| | Skin | Severe multifocal ulcerative dermatitis with intra-epidermal pustules, intra-epidermal vesicles, ballooning degeneration of keratinocytes, bacterial colonization, neutrophilic dermatitis, vasculitis, and panniculitis. | Moderate to severe multifocal ulcerative, neutrophilic, lymphoplasmacytic, eosinophilic, and histiocytic perivascular dermatitis, dermal vasculitis with neutrophils and multinucleate giant cells with mild perivascular edema, epidermal ballooning degeneration and vacuolization, and panniculitis. | Mild multifocal ulcerative dermatitis with fibrinosuppurative epidermal crusting, multiple intra-epidermal pustules, marked epithelial ballooning degeneration, focal eosinophilic intracellular inclusions, single cell acantholysis, focal degeneration of the dermal collagen, and mild to moderate multifocal neutrophilic dermatitis, vasculitis, perivasculitis, and panniculitis. |
| | Heart | Severe hypertrophic cardiomyopathy. | Severe hypertrophic cardiomyopathy with minimal focal lymphocytic epicarditis. | Moderate multifocal degenerative and fibrosing cardiomyopathy |
| | Lungs | Bilateral moderate multifocal interstitial mineralization, mild multifocal anthracosis, mild perivascular hemosiderosis, mild BALT hyperplasia, mild to moderate peripheral emphysema, and focal Langham's giant cell formation. | Bilateral mild to moderate peripheral alveolar emphysema with alveolar histiocytosis, intravascular neutrophilia with perivascular lymphoplasmacytic inflammation, and minimal to mild multifocal BALT hyperplasia with focal of type II pneumocyte hyperplasia and mild multifocal anthracosis. | Marked vascular congestion, marked multifocal to diffuse pulmonary edema, hyperinflation of peripheral alveoli, alveolar rupture, emphysematous alveolar bullae, minimal multifocal alveolar histiocytosis, and moderate focally diffuse pleural fibrosis |
| | Inguinal lymph node | Sinus histiocytosis with lymphoid hyperplasia, hemosiderosis, and erythrophagocytosis. | Moderate diffuse sinus histiocytosis with mild lymphoid depletion. | Mild diffuse lymphoid depletion and sinusoidal erythrophagocytosis and minimal multifocal ink accumulation. |
| | Axillary lymph node | Sinus histiocytosis with lymphoid hyperplasia, hemosiderosis, and erythrophagocytosis. | Mild lymphoid depletion. | There are no significant histologic lesions in the tissue sections examined. |
| | Placenta | Mild multifocal neutrophilic villitis with necrosis, vacuolization and fusion of syncytiotrophoblasts, multifocal necrosis and hemorrhage of the basal trophoblastic shell, syncytial knot formation and neutrophilic and lymphocytic intervillositis, and chorionic vessel vasculitis. | Mild to moderate multifocal neutrophilic and histiocytic intervillositis, multifocal ischemia, and focally extensive subacute parenchymal hemorrhge, and Minimal multifocal peri-villous fibrin. | Marked multifocal acute and subacute retroplacental hemorrhage, acute neutrophilic inter-villositis, syncytiotrophoblast degeneration, & syncytial knots. |
| | Decidua basalis | Mild to moderate neutrophilic deciduitis, mild diffuse hemosiderosis, and spiral artery endovascular extravillous trophoblasts. | Mild to moderate neutrophilic deciduitis with hemorrhage, vascular hylanization, fibrin | Mild multifocal acute deciduitis with mild multifocal necrosis and mild multifocal gross hemorrhage. |

| | | | | |
|---|---|---|---|---|
| | | | thrombi, multifocal ischemic necrosis, and hemosiderosis. | |
| | Spleen | Mild diffuse neutrophilic splenitis. | Mild diffuse hemosiderosis with moderate lymphoid hyperplasia. | Mild lymphoid depletion with marked red pulp congestion, sinusoidal histiocytosis with erythrophagocytosis. |
| **Fetus** | Final summary comments | The vascular congestion and autolysis are consistent with fetal death in utero. There is no evidence of inflammation.<br><br>There is no evidence of: overlapping sutures, prominent occipital bone, redundant scalp skin, ventriculomegaly, cerebellar hypoplasia, microphthalmia, coloboma, clubfoot, or contractures/arthryogryposis in this fetus. | The vascular congestion and coagulative necrosis in the liver are both consistent with fetal death in utero. This is supported by the findings in the umbilical cord in the Dam's necropsy report.<br><br>There is no evidence of: overlapping sutures, prominent occipital bone, redundant scalp skin, ventriculomegaly, cerebellar hypoplasia, microphthalmia, coloboma, clubfoot, or contractures/arthryogryposis in this fetus. | There is no evidence of autolysis or inflammation in any of the tissue sections examined. The focal areas of extravasation (hemorrhage) are most likely due to collection of cerebral spinal fluid and neuropil. The EMH in the liver is appropriate for the fetal age.<br><br>There is no evidence of: overlapping sutures, prominent occipital bone, redundant scalp skin, ventriculomegaly, cerebellar hypoplasia, microphthalmia, coloboma, clubfoot, or contractures/arthryogryposis in this fetus. |
| | Total body | Mild congestion and autolysis. | Mild to moderate diffuse autolysis, marked dermal edema with mottling. | No autolysis or edema. |
| | Skull & brain | Moderate to marked dermal capillary and muscular congestion with mild artifactual neural fragmentation. | Marked capillary congestion. | Mild congestion and focal hemorrhage in meninges. Mild diffuse congestion in choroid plexus. |
| | Liver | Mild to moderate autolysis with age appropriate extramedullary hematopoiesis. | Mild to moderate multifocal midzonal hepatocellular coagulative necrosis with multifocal hemorrhage and age-appropriate extramedullary hematopoiesis. | Moderate diffuse extramedullary hematopoiesis. |

**Table S1. Table showing results of qualitative histopathological analysis of maternal and fetal tissues.**

| Reagent | Manufacturer | Product number | Dilution | Assays in which reagent was used |
|---|---|---|---|---|
| Rabbit anti Vaccinia | Genetex | GTX36578 | 1:1000 (IHC) or 1:1500 (IF) | IHC primary, IF primary |
| Mach 2 Rabbit HRP-polymer | Biocare | RHRP520H | Neat | IHC secondary |
| CD163 | Genetex | GTX42365 | 1:100 | IF primary |
| Cytokeratin | Sigma | 452M-94 | 1:75 | IF primary |
| CD31 | Bioss | BSM10825M | 1:200 | IF primary |
| Goat anti rabbit IgG (H+L) Alexa fluor 647 | Invitrogen | a32728 | 1:1000 | IF secondary |
| Goat anti rabbit IgG (H+L) Alexa fluor 594 | Invitrogen | a32740 | 1:1000 | IF secondary |
| Rabbit IgG isotype control | Invitrogen | 31236 | 1:4000 | Control = IHC for vaccinia and IF for vaccinia |
| Mouse IgG2 | Santa Cruz | 3878 | 1:100 | Control = IF for cytokeratin |
| Mouse IgG1 | Santa Cruz | 3877 | 1:100 | Control = IF for CD163 and CD31 |

**Table S2. Table of reagents used in immunohistochemistry (IHC) and immunofluorescence (IF) experiments.**
